# Demographic Reconstruction of the Western Sheep Expansion from Whole-Genome sequences

**DOI:** 10.1101/2023.06.16.545369

**Authors:** Pedro Morell Miranda, André ER Soares, Torsten Günther

## Abstract

As one of the earliest livestock, Sheep (*Ovis aries*) were domesticated in the Fertile Crescent about 12000 *−* 10000 years ago and have a nearly worldwide distribution today. Most of our knowledge about the timing of their expansions stems from archaeological data but it is unclear how the genetic diversity of modern sheep fits with these dates. We used whole-genome sequencing data of 63 domestic breeds and their wild relatives, the Asiatic mouflon (*O. gmelini*), to explore the demographic history of sheep. On the global scale, our analysis revealed geographic structuring among breeds with unidirectional recent gene flow from domestics into Asiatic mouflons. We then selected four representative breeds from Spain, Morocco, the UK and Iran to build a comprehensive demographic model of the Western sheep expansion. We inferred a single domestication event around 9000 years ago, slightly later than archaeological evidence suggests which might reflect uncertainties in the generation time used for these estimates. The westward expansion is dated to approximately 5000 years ago, later than the original Neolithic expansion of sheep and approximately matching the Secondary Product Revolution associated with woolly sheep. We see some signals of recent gene flow from an ancestral population into Southern European breeds which could reflect admixture with feral European mouflon. Furthermore, our results indicate that many breeds experienced a reduction of their effective population size during the last centuries, probably associated with modern breed development. Our study provides insights into the complex demographic history of Western Eurasian sheep, highlighting interactions between breeds and their wild counterparts.

## Introduction

Sheep (*Ovis aries*) represent one of the earliest known livestock species to be domesticated in the Fertile Crescent about 12000 − 10000 years ago (1–4) and it has been a key resource for pastoral and agricultural communities ever since. Several studies have revealed aspects of sheep history and evolution that point to a complex demographic history with an uncertain origin and multiple waves of expansion from the Middle East/Central Asia into Europe. The presence of multiple distinct mitochondrial DNA haplogroups in modern sheep that were already present before domestication (5) has been suggested as evidence of multiple domestication events. However, a large and heterogeneous wild population, major gene flow from wild sheep into domestic flocks, or a combination of both could also explain this pattern. Studies on Y chromosomes (6) and polymorphic endogenous retroviruses (7) have suggested the possibility of important gene flow events after the initial expansion related with the development of wool industries.

Analyses of genome-wide datasets have been successful in providing important insights into the evolutionary history of sheep breeds across the globe. Studies using SNP arrays showed that the species exhibits general geographic clines that match the expansions into the different regions following the Neolithic transition (8–10). Contributions from diverse wild stock are also supported by the higher genetic diversity and haplotype sharing compared with other domesticates observed in modern breeds (8). At the same time, population structure and demographic history on a more regional scale are consistent with a scenario in which admixture with other sheep breeds and wild ovids is more common than previously thought (9–12) and may even have been encouraged in some cases to acquire desired traits from local wild ovids (13–15). More recently, whole genome sequencing has been employed for understanding global population structure and history (6, 14, 16), introgression from wild relatives (14– 16), how early artificial selection shaped domestic groups (17, 18), and patterns shared between sheep and other domesticate species (19). While these studies highlighted the power of whole genome sequencing data for insights into the past of this important livestock species, they were mostly focused on global patterns without addressing intra-continental patterns, aggregating together different breeds at the continental scale or by phenotypic characteristics (e.g. the presence of a fat or thin tail), which can lead to the miss-representation of continental or regional demographic patterns.

Today, sheep are a very popular livestock species along the Atlantic coast from NorthWestern Africa to the European Islands of the North Atlantic with hundreds of recognized breeds ranging from local landraces to popular industrial breeds. After their initial westward introduction into Europe, sheep have experienced at least one additional expansion from Western Asia, possibly associated with the development of wool industries (6, 7). In contrast to other parts of Eurasia, the absence of wild ovids in Europe facilitated feralization which later enabled back-admixture from long-term feral European mouflon into managed breeds (9, 15, 16). Europe was also the place where selective breeding as scientific practice started during the British Agricultural Revolution in the 18^th^ century (20). More recently, popular industrial breeds, such as Merino originating from Iberia, have been exported to other continents and were used for cross-breeding with many other breeds (21, 22). Altogether, this paints a picture of a complex demographic history of European sheep with a lot of uncertainties about the exact timing of particular events such as separation of different streams of ancestry or the presence and extent of gene flow post separation of populations and breeds. Whole genome resequencing data together with state-of-the-art statistical modeling approaches, however, should have the power to investigate these open questions.

The aim of this study is to infer the demographic history of Western Eurasian sheep from an extensive dataset of whole genome sequences, including both landraces and improved breeds as well as wild Asiatic mouflons (*O. gmelini*). We first performed an exploratory analysis on a global panel of sheep and mouflon in order to select five representative sheep populations and propose different demographic models for the history of Western sheep. These models differ in their general topology and some of them include admixture events. We estimate split dates as well as the extent of gene flow using the Site Frequency Spectrum (SFS) (23) and investigate how their effective population sizes changed over the last millennia and centuries. This approach allows us to describe the relationship between sheep breeds in Western Europe and how they were shaped by the demographic events of the past.

## Materials and Methods

### Data collection and processing

This study uses publicly available whole genomes from commercial and traditional domestic sheep breeds from the International Sheep Genome Consortium (ISGC) (24, 25) and a set of wild Asiatic mouflons (*O. gmelini*) from the *NEXTGEN* project (19). These datasets include 63 domestic sheep breeds from 21 countries and 935 individuals, and 18 wild mouflons from Iran (Table S1). In addition to whole genome data we gathered a set of 79 mitochondrial genomes from domestic sheep, urial (*O. vignei*), argali (*O. ammon*), snow sheep (*O. nivicola*) and bighorn sheep (*O. canadensis*), publicly available in GeneBank, which were then combined with the mitochondrial genomes from the wild Asiatic mouflons and 2 ancient samples (Table S4) from the Anatolian Neolithic, dated to 7031 − 6687 cal BCE and 6469 − 6361 cal BCE (26).

The 18 Asiatic mouflon genomes in FASTQ format were processed following the ISGC pipeline (25) to keep consistency with the already processed ISCG v2 dataset. FASTQ files were mapped to the Oar3.1 reference genome using *BWA mem* (v0.7.12) (27) with default parameters, and duplicates were removed with *SAMtools* (v1.3) *rmdup* (28). Local indel realignment was performed using *GATK* (v3.4.46) *RealignerTargetCreator* (29). Variant calling was performed independently using *SAMtools mpileup* and *GATK UnifiedGenotyper* with default parameters, and the resulting VCF files were filtered to remove multiallelic variants, variants never observed on one of the strands, low quality (PHRED score *<* 20) and low mapping quality variants (PHRED score *<* 30), variants where coverage was *<* 10*x* across all samples and in cases where there were 2 variants closer than 3 bp apart, the one with the lowest quality was removed. Similarly, in the case of indels that were closer than 10 bps, the lowest quality one was removed, and variants within 5 bp of an indel were filtered. Then, an intersect of both files was created to produce the final VCF file using *GATK CombineVariants* and *SelectVariants*.

To avoid issues with over-representation of some breeds, the dataset was sub-sampled to, at most, 5 randomly selected samples per breed, and only SNPs with a minor allele frequency (MAF) of 0.05 and a genotype rate of 0.1. These filters resulted on a dataset of 176 samples (Table S2) and 16932388 SNPs.

Mitochondrial consensus sequences were called from FASTQ sequences with *MIA* (30), a reference-based iterative assembler, using the reference sequence for the Asiatic mouflon (NCBI Reference Sequence NC_026063). To ensure reliability on the bases called, a minimum of 10 unique molecules (10*x* coverage) was necessary for a consensus to be called for each position, plus a minimum map quality of 40 and two-thirds base agreement on each position. Any site that did not meet any of these quality parameters was called “N”. All sequences were then manually assessed and aligned using *MAFFT* (v7.407) (31).

### Exploratory population genetic analysis

In order to explore the relationship between the different breeds, we performed a Principal Component Analysis (PCA) using *Smart-PCA*, from the *EigenSoft* (v7.2.1) package (32, 33). To avoid the effect of linkage disequilibrium (LD), data was pruned using *Plink* (v1.90b4.9)’s parameter `-indep` option (500 1 1.066) (34). Further population structure analysis was performed using *ADMIXTURE* (v1.3) (35). The number of assumed clusters ranged from 2 to 10, and each cluster was run with different random seeds 20 times. The results were compared and plotted using *Pong* (v1.4.9) (36).

Data was then sub-sampled to a set of 5 representative populations: Border Leicester and Merino as proxy for North-Western and South-Western European sheep, respectively, D’Man for North African sheep, and a group of Iranian sheep of no formally described breed, in the original data labeled as “Unknown”, for Middle Eastern sheep (Figure 1B, Table S3). To explore the splits and admixture events between our groups we created admixture graphs with *OrientAGraph* (v1.0) (37) using the LD pruned dataset and the *-mlno - allmigs*. To assess the robustness of the results and avoid local optima, *OrientAGraph* was run using the *-bootstrap - k* 500 option to create 10 replicates. All trees were visually inspected using the *Treemix* (38) plotting functions on *R* (v4.2.2) (39) and the one with the highest likelihood was then selected.

**Fig. 1.**
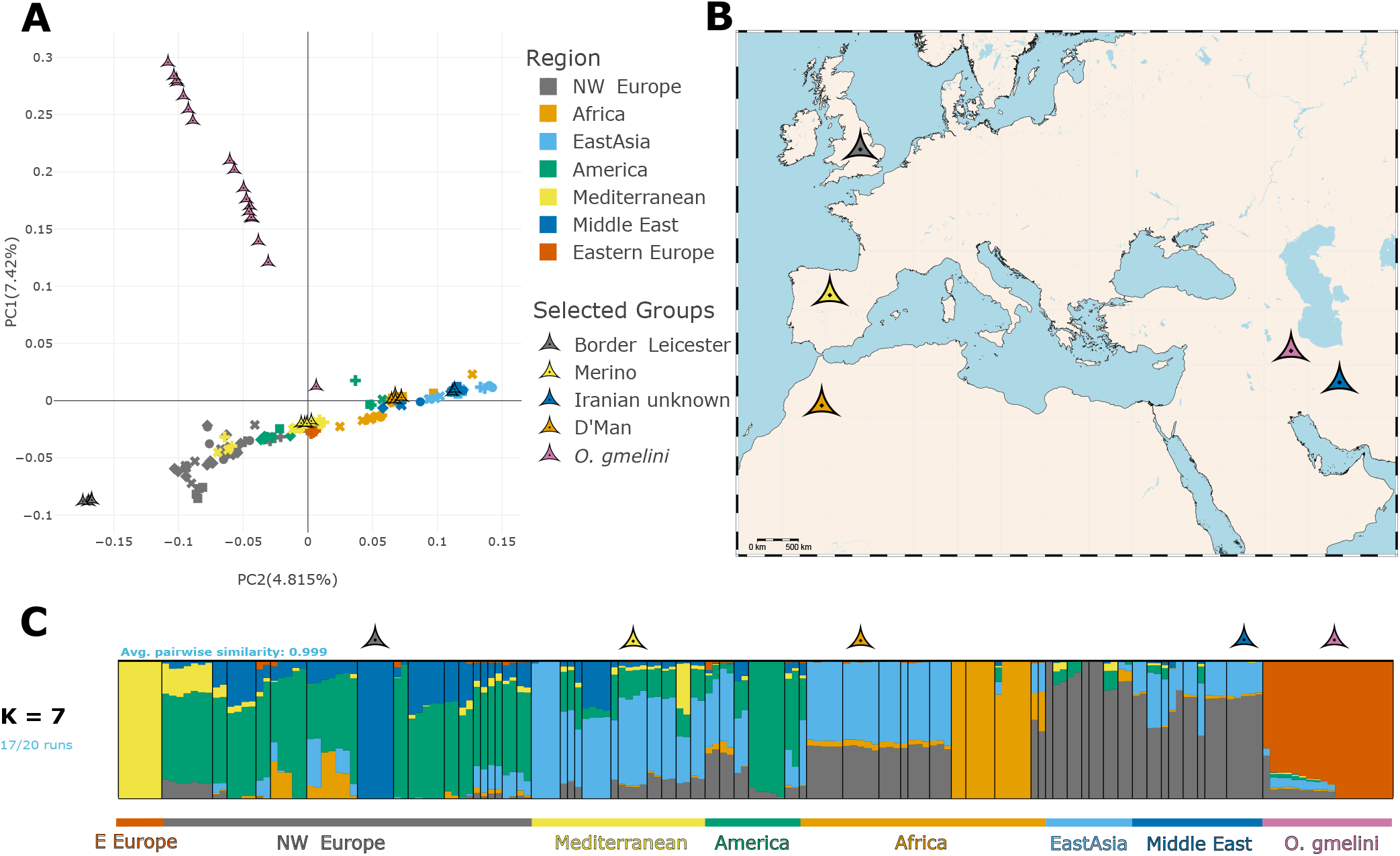
A) Graphic representation of the first two principal components (PCs) of a PCA calculated using 63 domestic breeds and Asiatic mouflons. PC1 describes the variation within wild mouflons, with some individuals showing signs of admixture with domestics sheep. PC2 captures the variation within domestic sheep and presents a geographical pattern, with European breeds on the negative extreme, and East Asian breeds on the positive. B) Geographical origin of the 5 breeds used for demographic inference. C) Proportion of each sample’s genome assigned to K=7 genetic clusters using *ADMIXTURE*. Wild mouflon ancestry is almost exclusively observed in wild samples (orange), while Eastern Eurasian samples present mostly a single ancestry cluster (in Grey). Middle Eastern and African samples show variable levels of ancestry from Eastern and Mediterranean ancestry (grey and light blue, respectively), and Mediterranean and North-Western breeds show high levels of admixture from each other (light blue for Mediterranean, dark blue and green for North-Western) and Eastern European (bright yellow).

### Demographic modeling

Informed by the results of the exploratory analysis, we defined five models for the demographic history of Western sheep to be tested with *Momi2* (v2.1.16) (23). The models were defined as follows:

- **Model A**: Single domestication event without admixture, assuming Iranian and Moroccan sheep form a monophyletic group.
- **Model B**: Single domestication event assuming Moroccan sheep are a sister group to European sheep. A variant of this model with admixture from wild mouflons into Merino, as suggested by *OrientAGraph* with 1 migration edge, was also considered.
- **Model C**: Single domestication event assuming Moroccan sheep are a sister group to European sheep with an admixture event into the Merino sheep, as suggested by *OrientAGraph* with 1 migration edge. In contrast to the gene flow variant of Model B, the source of the gene flow originates from an ancestral domestic ghost population and not the wild mouflon branch.
- **Model D**: Two independent domestication events from different regions inside of the Fertile Crescent for Eastern and European Sheep. A variant of this model with admixture from wild mouflons into Merino, as suggested by *OrientAGraph* with 1 migration edge, was also considered.
- **Model E**: Two independent domestication events from different regions inside of the Fertile Crescent for Eastern and European Sheep with admixture into Merino sheep, as suggested by *OrientAGraph* with 1 migration edge. In contrast to the gene flow variant of Model D, the source of the gene flow originates from an ancestral domestic ghost population and not the wild mouflon branch.

A schematic representation of all model topologies is shown in Figure 2. All branches in the models were allowed to have a different population size and all domestic breeds were allowed to grow freely, split times were only constrained by the order of splits defined by the model topology and with a starting value based on prior information (e.g. for the domestication time all models were set to start at 12000, see Table S7 for a detailed description). The initial mutation rate was set to 2.5 × 10^*−*8^ and the generation time to 2 (19). Allele frequencies and the SFS were calculated for 5 individuals of each of the 5 selected populations using *Momi2* (23), the optimization of the model was done using the *L-BFGS-B* algorithm and model fitness was evaluated both with log-likelihood and the Kullback-Liebler Divergence parameter (40). Confidence intervals (CI) were calculated using the results of *Momi2* bootstrapping runs (*n* = 100) for each parameter and then using *numpy*.*percentile* (41) with a confidence of 95%, and correlation coefficients were calculated using *scipy*.*stats’ spearmanr* function with standard settings.

**Fig. 2.**
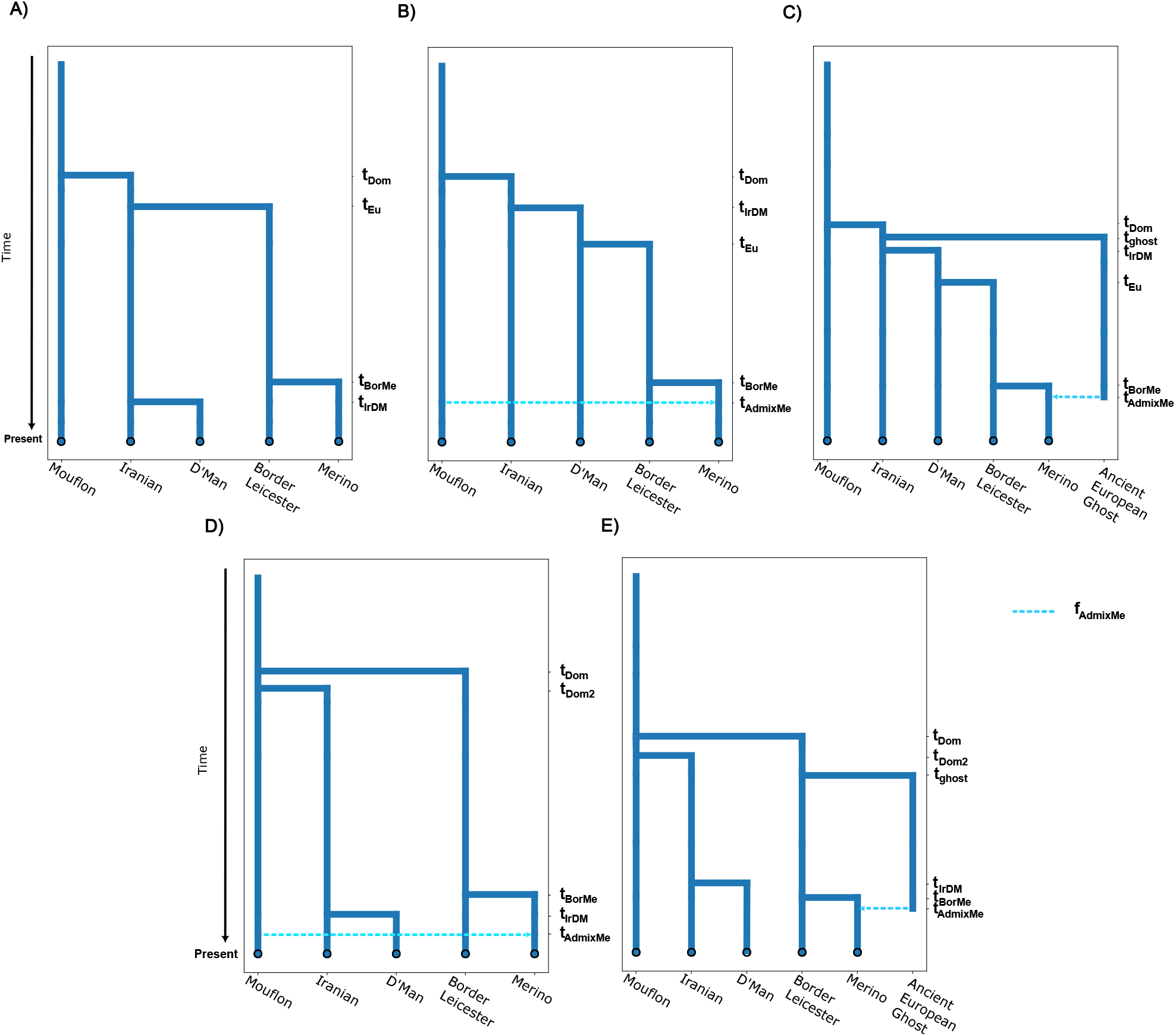
Topologies for the demographic models tested, displayed here with one admixture pulse (if admixture was considered). A) Single domestication event with Moroccan and Iranian sheep as a monophyletic group, B) Single domestication event assuming Moroccan sheep are a sister group to European breeds with an admixture event from wild relatives, C) Single domestication event assuming Moroccan sheep are a sister group to European breeds with an admixture event from a domestic ghost population, D) Independent domestication of Eastern and Western sheep with gene-flow from wild mouflons into Merino, E) Independent domestication of Eastern and Western with admixture from a domestic ghost population. The dashed lines represent an admixture pulse.

**Fig. 3.**
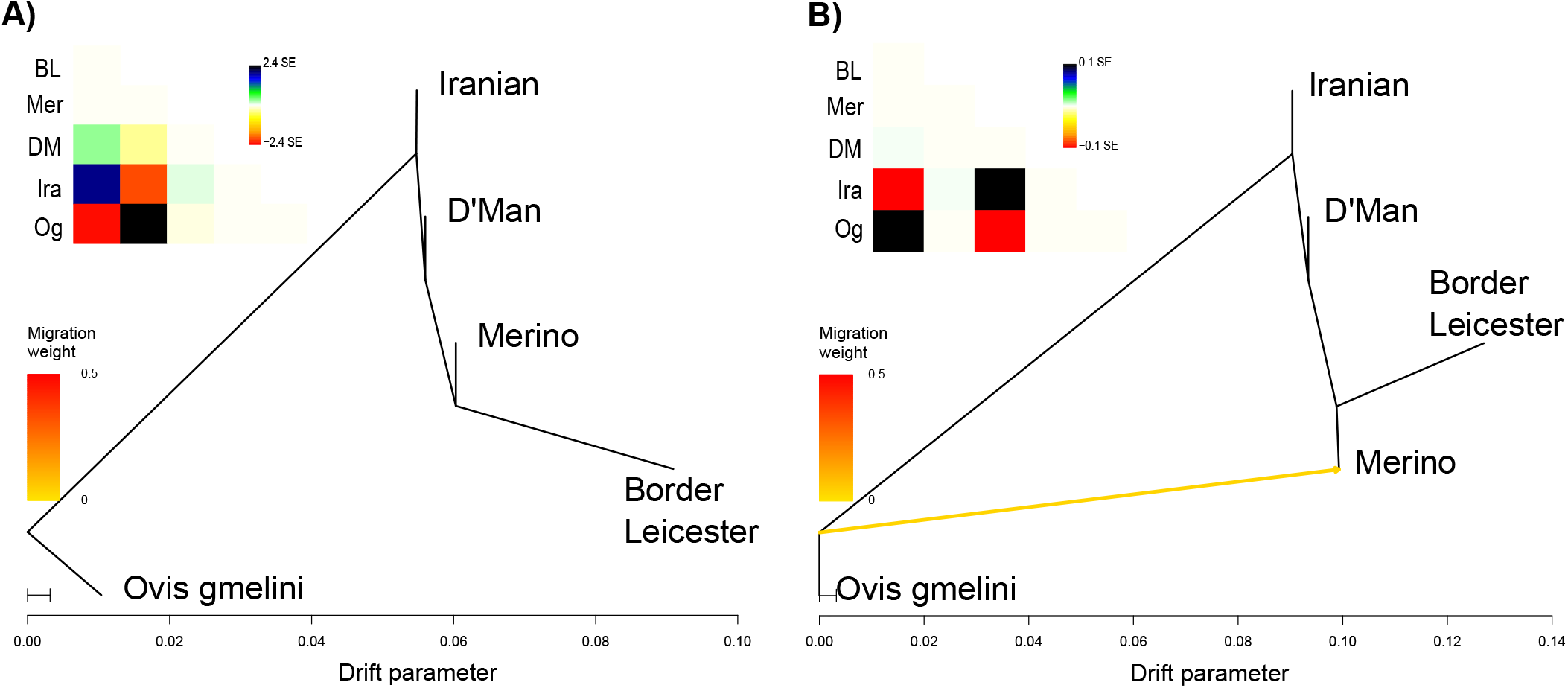
Maximum Likelihood Network Orientation reconstruction for the relationship between 4 domestic breeds and the Asiatic mouflon using *OrientAGraph*. A) The tree without migration edges displays a pectinate topology with the Iranian sheep splitting first, and D’Man being a sister group to both European breeds. The residual plot show high values for the intersection between wild mouflons and Merinos and between Iranian sheep and Border Leicester. These are resolved adding B) 1 migration edge from the base of the tree into Merino. The columns of the residual matrices follows the same order as the rows.

**Fig. 4.**
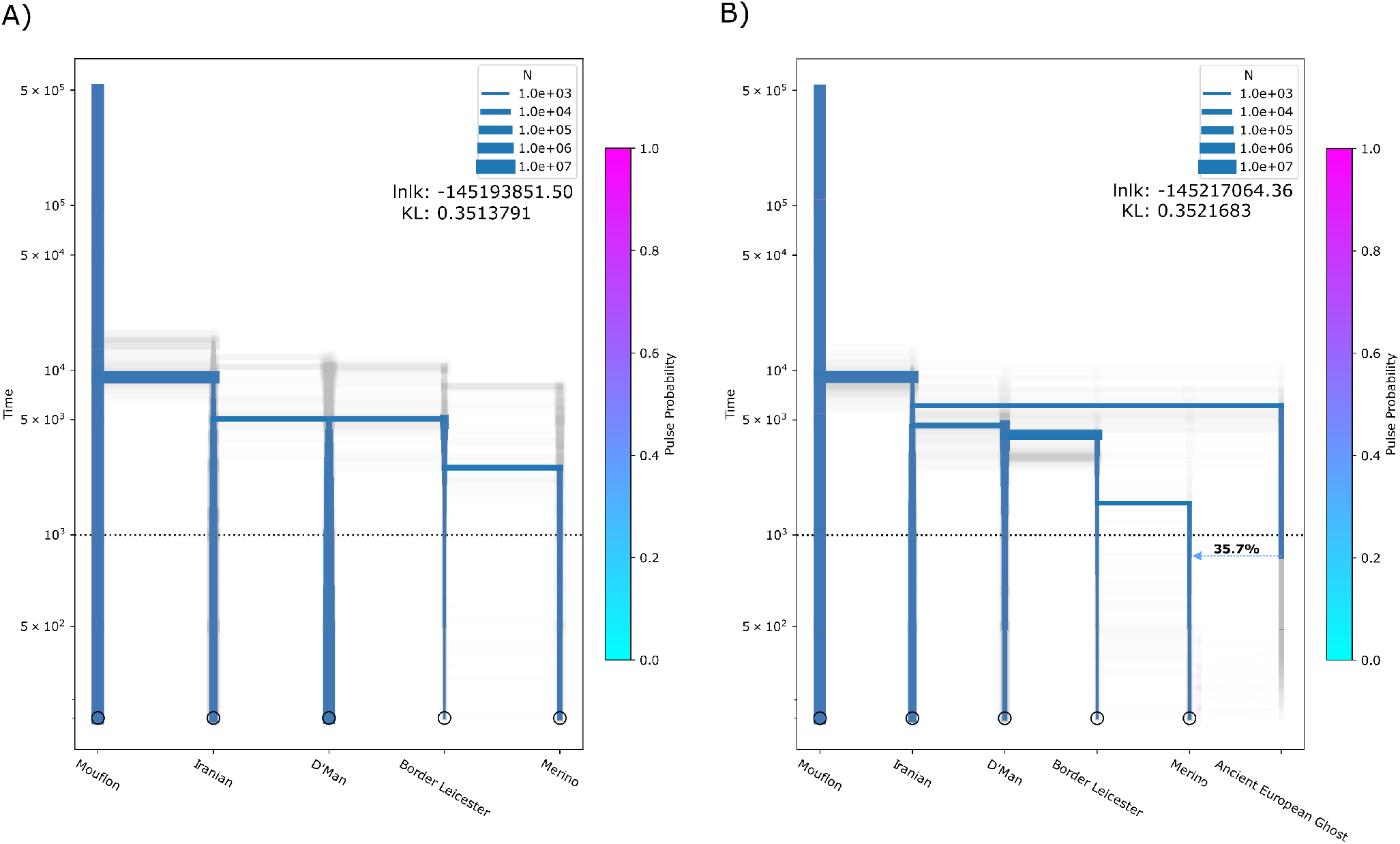
Maximum Likelihood inference of the demographic history of Western sheep using *Momi2*. A) The most likely model is Model B without admixture, and shows the same topology as *OrientAGraph* without migration pulses, with narrow confidence intervals around the expected time of domestication. B) The second most supported model is Model C, which displays the same topology as Model B, but adds admixture from a basal domestic ghost population. The model shows similarly narrow split times close to the expected domestication period, but shows bigger confidence intervals when dating the split between European breeds. Time is in years assuming a generation time of 2 years.

**Fig. 5.**
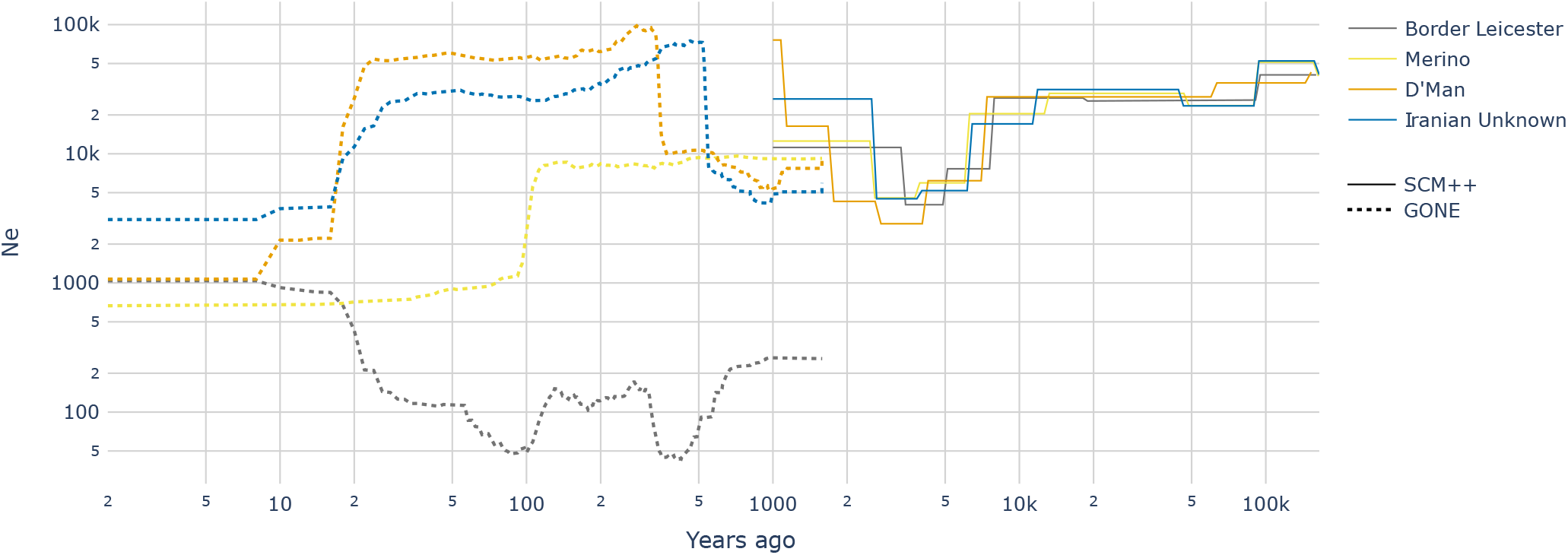
Effective population size inference of the four domestic sheep populations used in the demographic history reconstruction using *SMC++* and *GONE. SMC++* describes an initial bottleneck that starts around 10000 years, coinciding with the expected domestication period, with most breeds reaching a minimum around 3000 *−* 5000 years ago and a subsequent increase in N_e_ that is independent for each population. *GONE* describes more recent changes in N_e_. D’Man and the Iranian Unknown sheep seem to have experienced an increase on N_e_ around 500 and 300 years ago and then a bottleneck, while Merinos show a strong bottleneck 100 years ago. Border Leicester seems to have undergone several strong bottlenecks recently with a fairly recent rise in N_e_. Time is in years assuming a generation time of 2 years.

### Effective population size estimation

To investigate the effective population size (N_e_), we combined the results of two methods of N_e_ inference that do not overlap in time-span covered: *SMC++* (42) for deep-time N_e_ estimations, and *GONE* (43) for more recent changes. Data from all available samples from each population was used. *SMC++* input data was prepared using the *vcf2smc* function with default settings on all unphased autosomal chromosomes, and run with *estimate* with the same mutation rate per generation as *Momi2* models. *GONE* input data was prepared using *Plink* (v1.90b4.9) to produce *ped* and *map* files on unphased autosomal chromosomes and was run with standard settings except for a MAF filter of 0.1.

### Mitochondrial phylogeny

A phylogenetic analysis using mitochondrial data was performed. All sequences were aligned using *MAFFT* (v7.407) (31). We then created phylogenetic trees using both a Maximum Likelihood (ML) and a Bayesian approach (Bayesian inference, BI). For the ML tree, we used *IQ-TREE* (v2.1.3) (44), and allowed the algorithm to infer the best substitution model using *Model Finder Plus*. We estimated the phylogeny and divergence time of the alignment using the Bayesian genealogical inference package *BEAST* (v2.5) (45). We assumed the *GTR* + Γ nucleotide substitution model under an uncorrelated lognormal relaxed clock with a Yule process tree prior and two calibrations points (46). We used as calibration points the separation of *O. dalli* and *O. canadensis* from the Eurasian species (normal prior centered at 1.57 Mya, according to (47)), and the group containing *O. orientalis, O. ammon* and *O. vignei* (normal prior centered at 1.72 Mya (47)). We also used tip dating calibration for both Anatolian Neolithic samples (*tps062* and *tps083*) according to radiocarbon dates reported in (26). We combined three independent Markov Chain Monte Carlo (MCMC) runs to ensure proper mixing of the chain. Each chain ran for 100 million iterations, discarding the first 20% as burn-in. We visualized convergence of the MCMC chains by eye using *Tracer* (v1.6) (48) and calculated the maximum clade credibility tree using *TreeAnnotator* (v2.5) (49). The final tree was edited with *FigTree* (v1.4.3) (50).

## Results

The main goal of this study was to reconstruct the demographic history of the Western sheep expansion. The enormous number of possible models does not allow us to perform explicit demographic modelling for all possible breeds together and we need to restrict the model space by reducing the number of populations. Therefore, we initially performed an exploratory analysis of publicly available genome data from a worldwide data set of sheep breeds and wild Asiatic mouflon (19, 24). Based on the results of this analysis, we selected representative breeds for a model-based reconstruction of their demographic history from whole-genome sequences which is complemented by a phylogenetic reconstruction of the maternally inherited mitochondrial genome.

### Global population structure

To obtain a general overview of the relationship between sheep breeds of different regions of the world we performed a PCA of the genome wide variation. The first major axis of variation (PC1) separates wild mouflon from domestic sheep breeds (Figure 1A). Mouflons are spread along this axis, which may suggest some heterogeneity within this group with regards to the relationship to sheep breeds and/or gene flow between domesticated sheep and wild mouflon (6, 10, 51). PC2 shows a clear distinction between different domestic sheep groups, which exhibit a geographic pattern across Eurasia with North-Western European sheep showing more positive values while Eastern Eurasian breeds tend to negative values, and Mediterranean, Middle Eastern and African breeds are distributed along this East-West gradient. Breeds with a known history of admixture such as Romanov and Dorper sheep fall between the two big continental groups. These two modern breeds are known to carry mixed Eastern and Western ancestry and were bred to adapt to the harsh climate conditions in Russia and South Africa, respectively (52, 53).

The genomic clustering analysis with *ADMIXTURE* (35) presented a similar pattern (Figure 6) as the PCA, with *K* = 2 discriminating between Asiatic mouflon and domestic sheep while some mouflons show signals of domestic sheep admixture and a small proportion of mouflon ancestry is seen in several domestic sheep. The latter signal, however, disappears for *K* = 3 and above (Figure 6) where we observe a split between Eastern and Western sheep clusters with most commercial Western breeds heavily admixed. At *K* = 7 (Figure 1C), we see a complex pattern of regional ancestries, with Eastern Asian breeds showing low levels of cluster diversity, while Middle Eastern and African breeds show a higher component in common with Mediterranean sheep, and European breeds seem to be a mix of three clusters: one more common in Mediterranean and North African breeds and two mostly present in North-Western European breeds, plus some from gene-flow from Eastern Europe and Eastern Asia. Higher values of *K* produce various different modes according to pong, indicating multiple local optima (Figure 6). Across all *Ks*, some Asiatic mouflons show admixture from domestic sheep while gene flow from wild mouflons into domestic sheep (including the sympatric Iranian sheep) appears to be minor or non-existent.

**Fig. 6.**
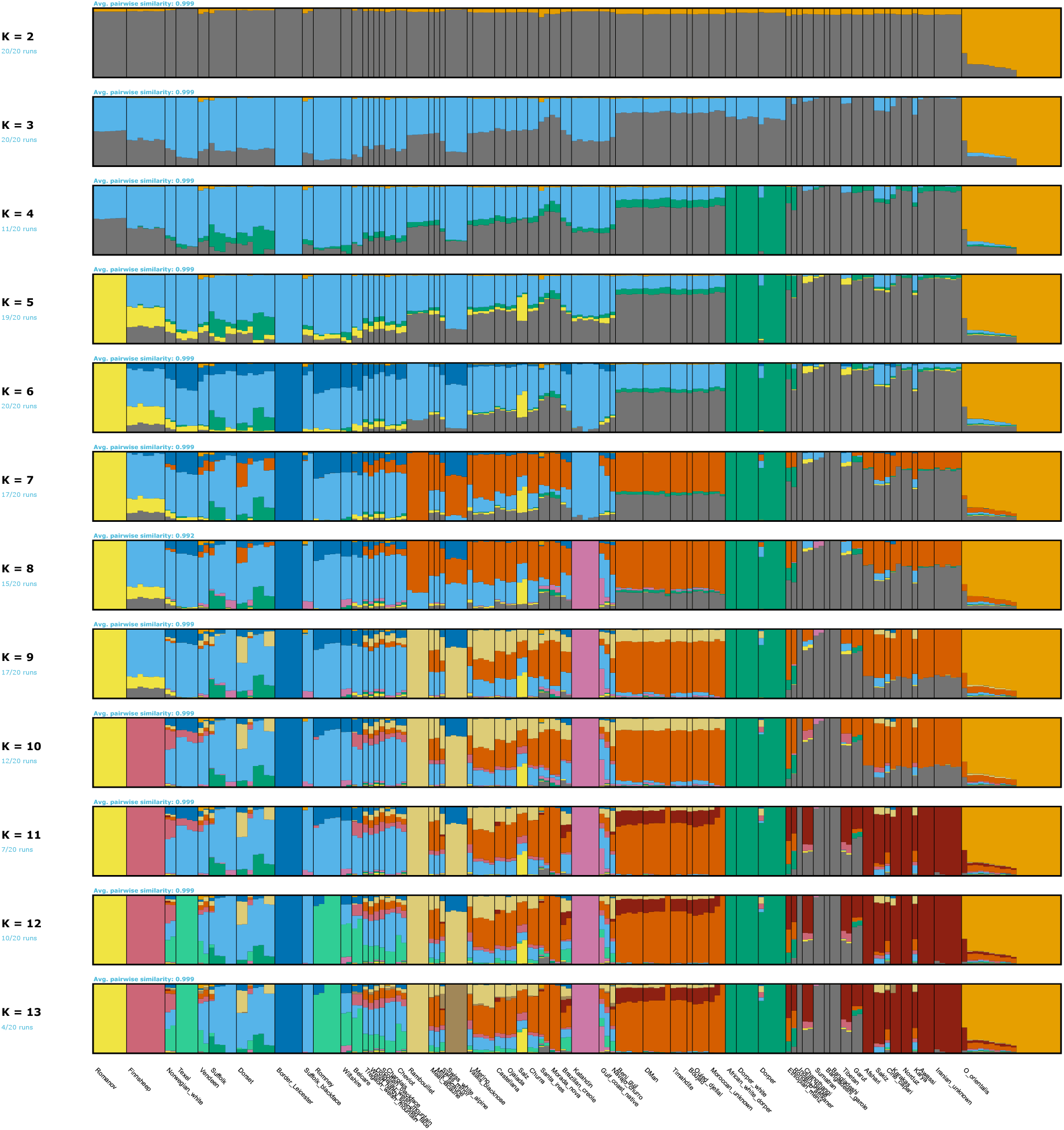
*ADMIXTURE* plots for 2 *−* 13*Ks*.

**Fig. 7.**
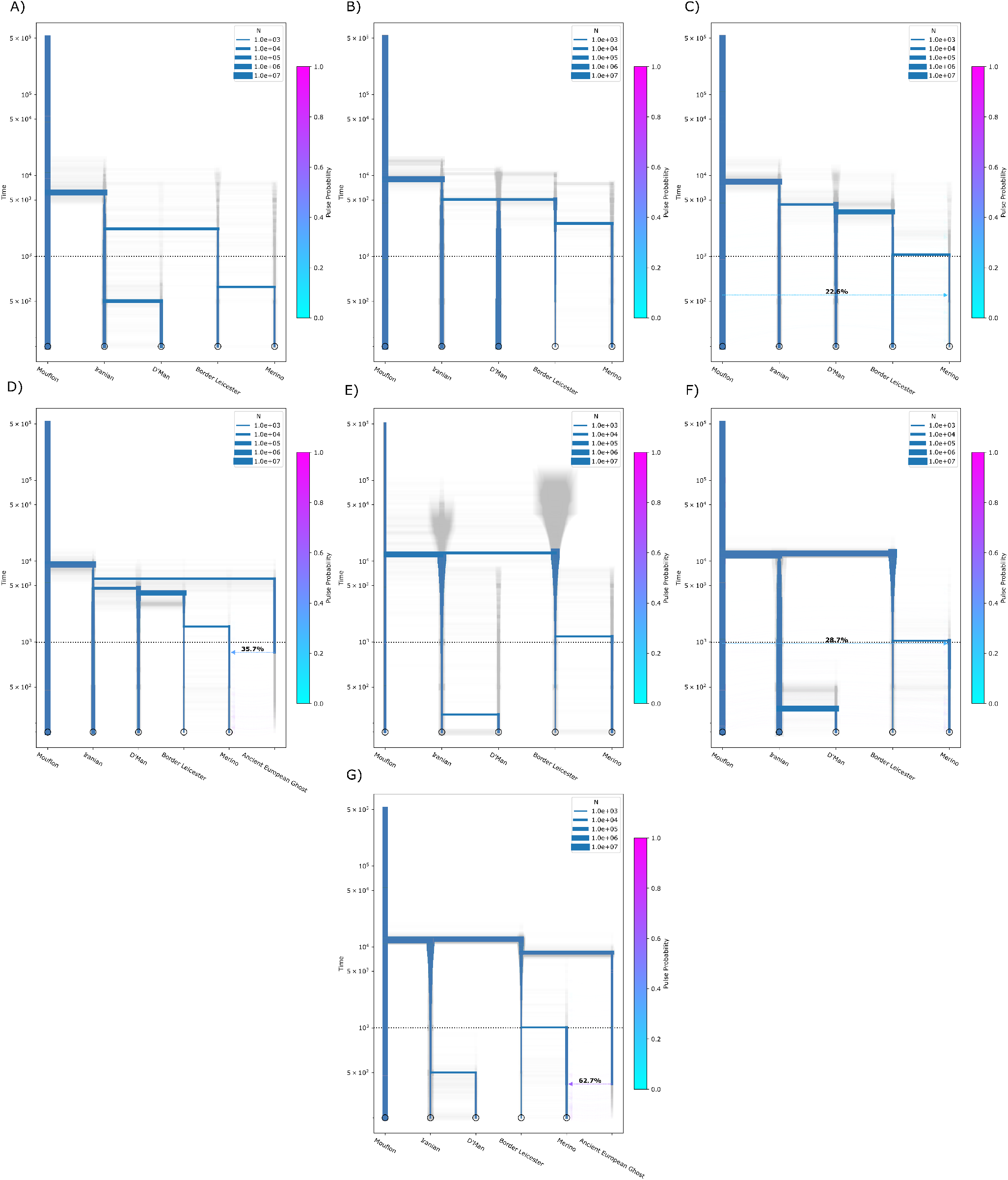
Results of *Momi2* demographic inference.

**Fig. 8.**
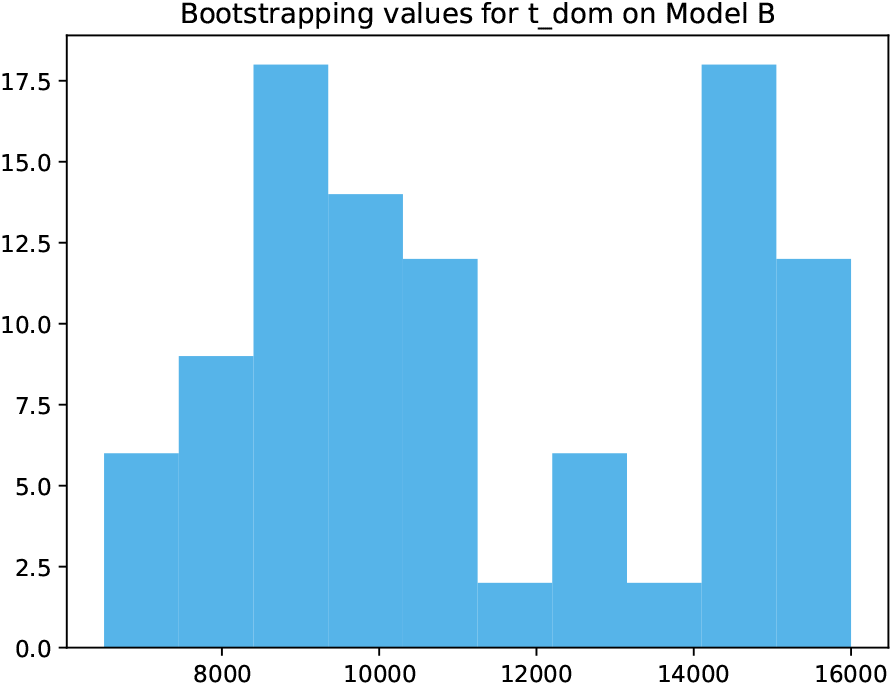
Bootstrapping values for *t*_*dom*_ on Model B. Most values cluster around the estimated value of model without resampling, but we observe a set of local optimum between 14000 and 16000 ya.

Gene-flow from from domestic species into mouflons is also supported by the mitochondrial phylogeny inference (Figures 9 and 10). Both trees show one mouflon clustering within the A haplogroup, with the most recent split being the Anatolian Neolithic sheep *tps083*. All other mouflons except one which seems to have been admixed with urials (*O. vignei*), belong to the CE haplogroup complex, but three mouflons also fall within a domestic branch, supporting the idea of domestic introgression from different sources into this wild population. Based on this exploratory analysis, we selected the wild mouflons as an outgroup and four domestic sheep populations covering different extremes of the Western sheep expansion for subsequent analysis (Figure 1B). The selected domestic populations were Border Leicester (United Kingdom), Horned Merino (Spain), D’Man (Morocco) and a set of traditional Iranian sheep that have not been properly defined into a breed, which are labeled as “Unknown” in the original data. These breeds were selected to serve as proxies for the origin and the extremes of the Western expansion of sheep from the Fertile Crescent, they had a sufficient number of samples for further analysis (*>* 19), and since both PCA and *ADMIXTURE* show them as independent populations from each other. More comprehensive analyses of the global population structure of sheep have been previously published (8, 10, 14, 17, 18).

**Fig. 9.**
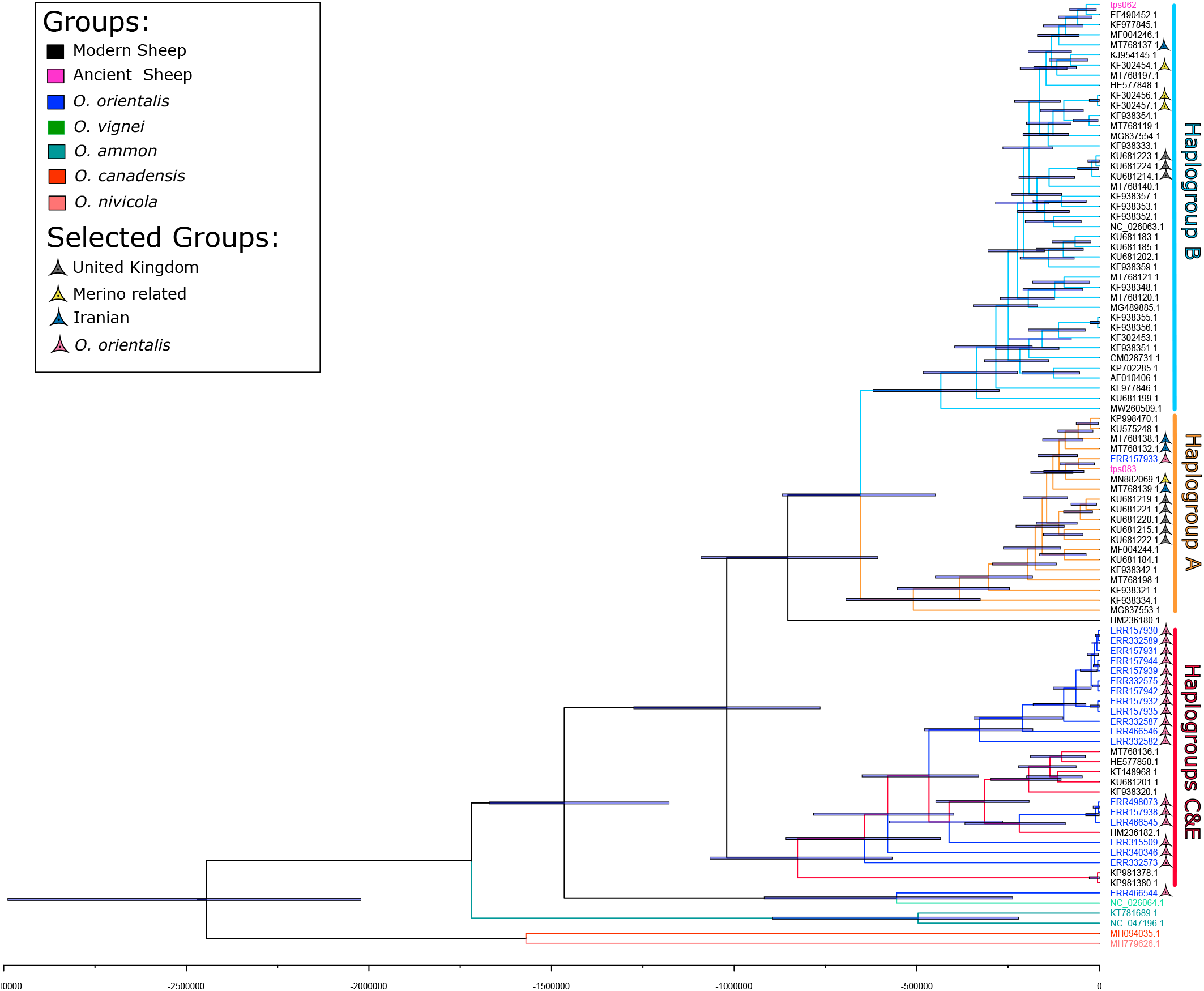
Bayesian Phylogenetic inference tree with *BEASTt*. It shows domestic sheep clustering into three groups matching the mitochondrial haplogroups A, B and the CE complex. Split time estimation between the different mitochondrial sheep lineages and the results corroborate previous studies (5, 6, 62, 63), which found that the three major haplogroups (A, B and the CE cluster) significantly predate the domestication of this species usually assumed to have taken place around 10000 years ago. Our estimates for the most recent common maternal ancestor between CE and (A, B) is 1020342 years ago (95% HPD: 764505 to 1274862) while the split between haplogroups A and B was dated to 653310 years ago (95% HPD: 448786 to 868374).

**Fig. 10.**
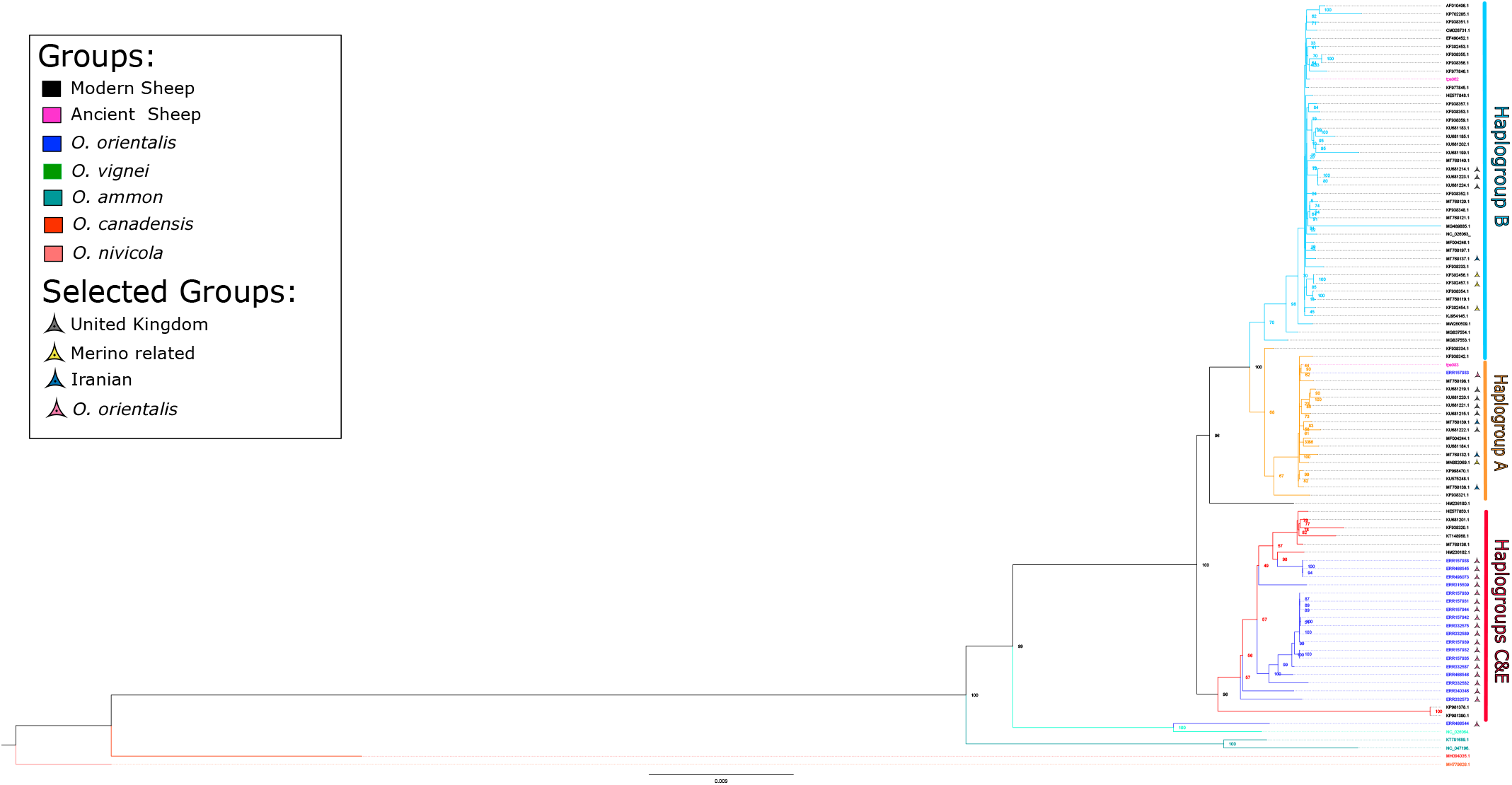
Maximum Likelihood consensus tree showing similar structure than *BEAST*, splitting into three clusters matching with Haplogroups A, B and the CE complex. Within-haplogroup resolutions, however, is lower, which several splits unresolved or with low support. Haplogroups A and B are ubiquitous across various Western sheep breeds making this analysis uninformative about the recent demographic history of the breeds discussed above. It does, however, provide additional insights into the relationship of the used mouflon populations to domestic sheep breeds: All but two Asiatic mouflons clustered within the CE complex. One mouflon falls within haplogroup A, close to the Neolithic *tps083*, while the last mouflon clusters with an urial (*O. vignei*).

### Structure and gene flow in Western Eurasian sheep

Using the representative subset of breeds as outlined above, we moved towards a more comprehensive analysis of their relationship. We used *OrientAGraph* (37) to explore possible topologies of models to test. The initial tree (without migration edges) follows a pectinate topology with Asiatic mouflon as an outgroup to domestic sheep then Iranian “Unknown” splitting off followed by D’man and finally Border Leicester and Merino as sister groups (Figure 3A). The residual covariance suggests some shared, unexplained ancestry between Asiatic mouflon and Merino which is resolved by a migration edge from the root of the tree into Merino (migration weight 0.0379) once one migration event is allowed (Figure 3B). At this point, most of the allele frequency covariance matrix is explained by the model with residuals only around ± 0.1 standard errors (SE). Therefore, we decided to restrict our models for the more explicit demographic reconstruction to models with maximum one gene flow event.

### Demographic reconstruction

Following these results and scenarios based on the literature (5, 9, 10), we defined five demographic models in order to reconstruct the demographic history of Western sheep (Figure 4). We used the SFS inferred from the whole genome sequences of the five populations to test these models using *Momi2* (Figure 7 and Tables S5 and S6) (23). The best-fitting model was model B (*logLikelihood* : − 145193851.50, Kullback-Leibler divergence, or *KLdiv*, between the predicted and observed SFS: 0.3513791), which assumes a single domestication event and that North African sheep are a sister group to European sheep and no admixture. Closely behind was Model C (*logLikelihood* : − 145217064.36, *KLdiv* : 0.3521683), which describes a similar topology, but with an admixture event from a basal European ghost population into Iberian sheep. Other topologies and Model B with admixture from wild mouflons into Iberia perform substantially worse. Models that assumed a double domestication were the ones that performed worst (Table 1). Lastly, if we compare the models that considered an admixture event, a domestic ghost population had always stronger support than the wild mouflon as the source of gene flow (Table 1).

**Table 1.**
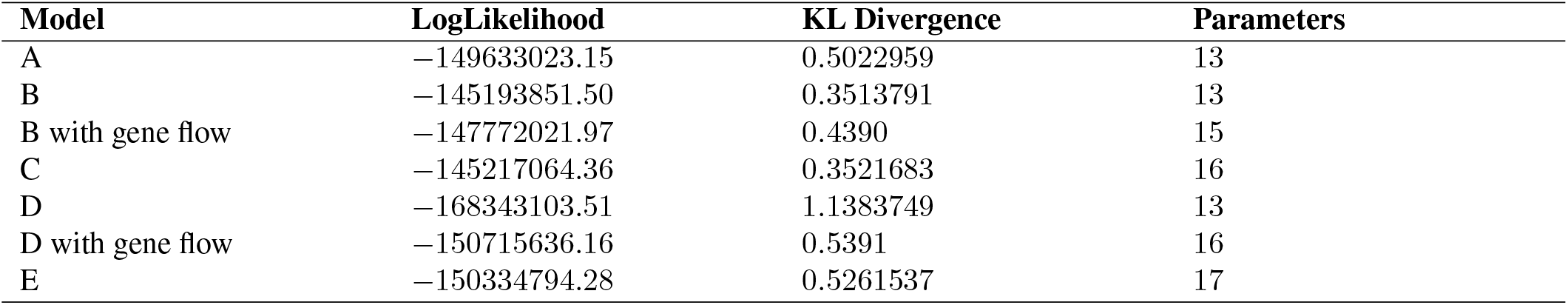
Performance of the different models tested by *Momi2:*

Estimated values for split times in the two best models roughly agree with archaeological evidence (54, 55). Model B estimates that the split between wild mouflons and domestic sheep happened 9062 years ago (ya) (95% CI [7006, 16000]). We also need to note that Model B displays a bimodal distribution of split dates across all bootstraps with the point estimate falling into the lower peak, a second set of local optima represents earlier split dates (initial split slightly over 14000 ya) (Figure 8). For Model C a very similar split time is estimated at 9103 years in the past (95% CI [6816, 14544]). Both models show that the split between Iranian and Western sheep and North African and European sheep happened in quick succession (5062 and 5058, respectively, for Model B, and 4636 and 4065 for Model C) (95% CIs [2603, 10500]; [3006, 12000]; [2558, 5283] and [2558, 8170]) (Table S6) but after the initial expansion of this species westwards during the Neolithic. Both models, however, disagree on the timing of the European split, with Model B estimating it at 2558 ya (95% CI [103, 8000]) and Model C at 1565 ya (95% CI [58, 2783]). This final split is also where both models’ bootstrapping values show an extreme spread (Figure 4), which seems to be in connection with the the admixture pulse in Model C estimated to have happened at 886 ya with a weight of 0.36 (Figure 4). These estimates also have quite wide CIs, ranging from 3 to 1614 ya and a weight between 0.08 to 0.9. However, those values are not randomly distributed, but negatively correlated. The more recent the event is estimated to be, the higher the weight (Spearman correlation of 0.4545, p-value of 2.6 × 10^*−*6^).

In terms of effective population size of our four domestic breeds, both models show a steep bottleneck in the population ancestral to all domestic breeds, as expected for this kind of process. The Iranian branch, however, quickly starts to grow in both models. Both models also agree that the ancestral population of the two European breeds starts shrinking shortly after the split from D’Man. The effective population size in Border Leicester continues decreasing while the size of the Merino branch is small but relatively steady. The two models show contrasting patterns for the demographic history of the Moroccan population, with Model B showing an initial bottleneck and later population growth, while Model C estimates a large initial population size and modest population size reduction since then.

### Effective population size over time

The *Momi2* results indicate a reduction of population size in each individual breed after they split from the other populations. To obtain a more detailed picture on this development with temporal resolution, we employed a combination of *SMC++* for the last millennia(42) and *GONE* for the last centuries (43) (Figure 5). *SMC++* shows a bottleneck beginning at the time of domestication (12000 − 8000 years ago) and after that initial decline each population increases in size at different times and rates in the last 3000 years. *GONE* estimates are considered most reliable in more recent periods (43). Indeed, we see that the *GONE’*s N_e_ estimates for about 300 − 500 years ago are very similar to the most recent estimates from *SMC++* from about 1000 years ago for all breeds except Border Leicester, which has a much lower population size according to *GONE* compared to *SMC++*. The other breeds show a reduction of their effective population size during the last centuries which could be associated with the appearance of modern breeding practices (56) and, in the case of Merino, the fact that the sequenced samples had been introduced to Australia. Border Leicester show an almost mirrored trajectory with extremely low effective population sizes up to the last century followed by an increase to similar levels as the other breeds. Border Leicester derives from Dishley Leicester, one of the first breeds subjected to selective breeding as a scientific practice (57) making it possible that the bottleneck associated with modern breeding practices started earlier and/or was more aggressive for these sheep. The recent recovery of effective population size could reflect the current popularity of the breed in Great Britain and other regions of the world, or could correspond to cross-breeding with other stocks although it is disputed whether Border Leicester were kept purebred during the last centuries (57). In both methods, consistent with the *Momi2* results, the final effective population size is lower in the two commercial European breeds.

## Discussion

### Structure of modern sheep populations and gene flow

Our analysis corroborated the known geographical structure of domestic sheep populations (8, 10, 26, 58). This geographical pattern is consistent with all current hypotheses about the origin and following expansion of sheep. *ADMIXTURE* results (see Figure 1C) suggest that admixture between domestic sheep breeds has been common, especially in Europe but also in Africa and the Middle East, where both commercial and traditional breeds display a variety of ancestry components both from Europe and other regions of Eurasia. In traditional breeds these seem to mainly receive admixture from geographically linked regions (e.g. Northern Africa populations being a mix of Mediterranean and Middle Eastern ancestries), with only minor contributions of really distant populations (e.g. small components of Northern European and Eastern European components in Eastern Asian breeds). In Europe, however, admixture between European breeds has been common, and Mediterranean breeds also show a significant amount of admixture with Eastern breeds. This pattern may be explained by constant trade and population movements during historic times, plus the advent of modern breeding practices in Europe and their expansion in later colonial times.

While we see this widespread gene flow between domestic groups, there is little evidence to support the hypothesis of major admixture of domestic sheep breeds with Asiatic mouflons. The only populations that show some level of wild mouflon ancestry are some European breeds, matching previously published results (19) (Figure 1). An alternative source for this signal is admixture with a population related to ancestral domestic sheep, as suggested by both *OrientAGraph* (Figure 3) and *Momi2* results (Figure 4). We propose that this archaic population is the European mouflon, assumed to represent a feral Neolithic sheep lineage from the islands of Sardinia and Corsica. Previous studies using low-density SNP genotyping suggested low levels of European mouflon admixture into Iberian sheep breeds (9, 10). Our second bestfitting model in *Momi2* is consistent with this interpretation as the timing of the admixture event into Merino ancestors would be extremely unlikely from an Asiatic mouflon. In addition to this, the inferred split date for the ghost population matched the expected date of feralization of the European mouflon (16). In contrast, gene flow in the opposite direction, from domestic sheep into wild Asiatic mouflon, appears to have taken place more frequently (Figure 1A and C, Figure 9, Figure 10).

### Demographic modelling and split dates

Our demographic reconstruction shows little to no support for the multiple domestication hypothesis. *OrientAGraph* and *Momi2* support a single domestication event with a pectinate topology (Figure 3 and Figure 4). In fact, when we fitted the double domestication models to our data, the estimated dates for both domestication events overlapped temporally at ∼ 12000 ya (best performing model’s 95% CI [11510, 14459]) (Table S6). Thus, they resemble a trifurcation soon after a single domestication event, but not necessarily two independent domestication events from different wild populations. We need to highlight, however, that our focus was the Western expansion of sheep and we only had sequences from an Eastern Asiatic mouflon population available (see also discussion below), which leaves possibilities for different contributions from other mouflon populations and a different history for Eastern Eurasian breeds.

While the exploratory analysis of the global dataset highlighted the abundance of recent admixture events between individual breeds, the motivation for the selection of the five populations used was to study deep demographic changes. Consequently, our demographic analysis points to little to no admixture between these five populations. Our two best fitting models’ split dates between wild mouflon and domestic sheep are just slightly over ∼ 9000 ya but with CIs ranging from 7006 to 16000 and from 6043 to 12160, respectively. The point estimate is surprising as it appears to post-date archaeological evidence for the domestication event (54, 55). Furthermore, the sequenced modern wild mouflon population is not likely to represent the direct descendants of the wild ancestors of domestic sheep which should push the estimated split times further back in time. While the historical range of this species completely covered the putative domestication regions, its current range only marginally overlaps with it (54, 59) suggesting that the original source population may no longer exist in the wild. Wild mouflon populations seem stable in their total range (16, 59) but their populations at a local scale have decreased in the last millennia (59, 60), which matches with their disappearance from regions of the Fertile Crescent (61). Furthermore, the genetic variation found in domestic sheep also does not appear to be a subset of the sequenced mouflon population. This is exemplified by the mitochondrial haplogroups, Asiatic mouflons clustered with the CE haplogroup, while most Western breeds fall into haplogroups A and B. The maternal split between CE and (A, B) is even dated to about one million years ago predating the domestication by two orders of magnitude consistent with other studies (5, 6, 62–64).

The recent estimates for the autosomal split may be explained by early gene flow from different wild populations after the initial domestication or by uncertainty regarding the appropriate generation time and mutation rates used to obtain these estimates. Our analysis used 2 years as generation time for sheep but assuming 3 years, as some studies did (14, 16, 18), would move Model B’s point estimate to ∼ 13500 ya. On the other hand, *SMC++* results yield credibility to a generation time of around or just above 2 years as the onset of the domestication bottleneck is dating to around or after 10000 ya (Figure 5). A recent study used simulations and a Maximum Likelihood approach to infer the split time for the onset of the domestication bottleneck, which was dated to 9078 using a generation time of 2 years (14). Using a generation time of 3 years, another recent study employed Approximate-Bayesian Computation to date the sheep/mouflon split to about 10500 years ago (6), similar to our estimates and closer to the estimated domestication period based on archaeological remains. The 95% highest posterior density interval (HPDI), however, included the maximum of the prior distribution used in that analysis (12000 years) suggesting that the point estimate of 10500 may be an underestimation for a generation time of 3 years. It appears that estimates based on a generation time of 2 years tend to post-date archaeological estimates suggesting that the historical generation time of sheep was slightly higher. In summary, the recent estimate for the split time is likely due to a combination between the uncertainties of the different estimates and a complex domestication process that is not well modelled by a simple split event.

The second split (∼ 5000 years ago) in our best supported model describes almost a trifurcation between the Iranian, North African and European branches, with only a few years of difference between population splits. This is several thousand years after the initial Neolithic Expansion, which means that this tree is not reflecting the initial expansion of sheep after their domestication. These split times, nonetheless, slightly predate the appearance of the Secondary Product Revolution and the presumed expansion of woolly sheep from Central Asia into the Fertile Crescent, Europe and Northern Africa (65, 66). As at least all the Western breeds display the woolly phenotype, this may be interpreted as most parts of the modern gene pool reflecting a secondary expansion westwards.

Both models had issues dating the split between Border Leicester and Merino, with CIs that range from over a century ago to the upper limit set for this parameter (8000 years) in the best performing model. Model C works slightly better when trying to date this split (95% CI [58, 2783]), but the estimate is 1000 years more recent and no clear local optima are observed among the bootstraps.

## Conclusions

Our results add to a mounting body of evidence on the complexity of the demographic history of sheep as well as the limitations of using genomic data from present-day populations to reconstruct it. Even for a restricted sample of five populations and a restricted geographic focus it remains difficult to narrow down certain demographic events. Recent demographic events in both commercial breeds and wild relatives make inferring ancient events particularly difficult, as the near-extinction of the Asiatic mouflon in a significant part of its original range or modern breeding practices could complicate demographic analysis. We were able to discriminate between several demographic models in favour of a single domestication event with domestic sheep radiating from the Fertile Crescent both eastwards and westwards. Dating these events, however, proved quite tricky. Our results point to the domestication happening from a highly diverse stock of Asiatic mouflon. This high diversity could be interpreted as evidence for multiple domestication events or, as our modelling suggest, a single domestication event in combination with introgression from diverse and significantly structured Asiatic mouflon populations, which we have not sampled or are not even present today. It is difficult to study such ancient events from the distribution of modern diversity alone and genomic ancient DNA from the domestication region could illuminate the demographic history of sheep as archaeogenomic studies have led to an enormous improvement in our understanding of domestication and subsequent gene flow in goats (67, 68), pigs (69) and cattle (70) already.

## Data availability

Data from the Sheep Genome Consortium v2 dataset can be accessed through the CSIRO data portal (https://doi.org/10.25919/5d39e494936c6), while the wild mouflons from the NextGen Project can be accessed via the project’s webpage in Ensembl (https://projects.ensembl.org/nextgen/). All mitogenomes were accessed via GeneBank and a list of accessions can be found in Table S4. The dataset used for the demographic inference and Momi2 models are available in Zenodo (https://doi.org/10.5281/zenodo.8017082). No new data was generated for this project.

## Acknowledgments

We thank the International Sheep Genomics Consortium as well as the NextGen and Sheep Genomes Database projects for making the sequenced genome data available to the research community.

## Supplementary Material

